# Effects of diet particle size on growth performance of the edible cricket, *Teleogryllus occipitalis* (Orthoptera: Gryllidae)

**DOI:** 10.1101/2024.04.22.590628

**Authors:** Kohyoh Murata, Takeshi Suzuki

**Affiliations:** Graduate School of Bio-Applications and Systems Engineering, Tokyo University of Agriculture and Technology, Koganei 2-24-16, Tokyo 184-8588, Japan

**Keywords:** cricket farming, digestion, feed design, field cricket, mandible

## Abstract

Farming edible crickets has environmental and nutritional benefits, as well as social benefits such as livelihood diversification. Commercial feeds for poultry and fish farming are often used to feed crickets, and in recent years, crop and food-processing by-products have also been used to improve sustainability. However, the design of feed for crickets has not been standardized. Here, we investigated growth and development of the Asian field cricket, *Teleogryllus occipitalis* (Audinet-Serville) (Orthoptera: Gryllidae), fed on different forms of the same diet. Body weights and the rate of development were significantly greater in crickets fed on millimetre-order granules than in crickets fed on micrometre-order powder. The result suggests that the granular form is easier for *T. occipitalis* to grasp and ingest than the powdery form, or that greater hydrophobicity of the powdery form inhibits digestion. Simply feeding millimetre-order granules may contribute to the development of feed design for farming edible crickets.

## Introduction

Edible-insect farming has attracted worldwide attention as an environmentally friendly source of animal protein and micronutrients (van Huis et al., 2013). Crickets are one of the most suitable insect groups for farming because of their rapid development, high fecundity, omnivorous nature, and ability to grow on dry feed as long as they are watered. In addition, the feed conversion ratio of crickets is 5.9 times that of beef cattle (van Huis, 2013), and the amount of greenhouse gases emitted by cricket farming is about one-quarter (CO_2_ + CO_2_-eq.) that emitted by beef cattle farming (Oonincx et al., 2010). The use of crickets as a food source is growing. In Thailand, approximately 20,000 farmers produce crickets, and annual production averaged about 7500 t from 1996 to 2011 (Hanboonsong et al., 2013). The European house cricket, *Acheta domesticus* (L.) (Orthoptera: Gryllidae), is the third insect species to be authorized as a novel food by the European Commission (2022).

Commercial feed for poultry and fish farming has often been used as cricket feed (Hanboonsong et al., 2013; Miech et al., 2016). In poultry farming, it is well known that the particle size of feed affects growth performance. For example, the weight of broilers fed a diet in crumb or pellet form tends to be higher than that of those fed mash, which is in powdered form (Reece et al., 1985; Svihus et al., 2004). Patton et al. (1967) described that *A. domesticus* nymphs tended to prefer smaller particles as diet to larger ones, but they presented no objective data such as particle size. No other studies have reported the effect of diet particle size on the performance of crickets.

Here, we show that diet particle size on its own affects nymphal survival, developmental time, and adult body weight of the Asian field cricket, *Teleogryllus occipitalis* (Audinet-Serville) (Orthoptera: Gryllidae), a traditional food species in East and South-East Asia.

## Materials and methods

### Insects

We used a population of *T. occipitalis* collected on Amami Ohshima (Kagoshima, Japan) and previously used for whole-genome sequencing (Kataoka et al., 2020). This population was maintained on chicken feed (Chougenki Edzukeyousuuyou; Nosan Corp., Kanagawa, Japan) or goldfish feed (Kingyo Genki Probio Flake; GEX Corp., Osaka, Japan) and water in polypropylene containers at 25 °C.

### Diet processing and particle size distribution

We bought an experimental diet for insects (I; Oriental Yeast Co., Ltd., Tokyo, Japan) in granular form. The powdered form was prepared from it in a food processor (SG-10BKJ; Conair Japan G. K., Tokyo, Japan). We took images of the granular diet with an image scanner (GT-X980; Seiko Epson Corp., Tokyo, Japan) and of the powdered diet with a scanning electron microscope (TM-3030; Hitachi High-Tech Corp., Tokyo, Japan). We used the images to calculate the particle size distribution by using the ROI manager and selection brush tool of NIH ImageJ 1.53 k software (Schneider et al., 2012) to measure the area (*S*) and calculating the particle size as that of a circle with the same diameter (Φ), as:

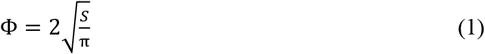

### Bioassay

A polypropylene container (239 mm × 176 mm × 91 mm) with a lid perforated with ∼100 holes (2.5 mm in diameter) for ventilation was used as a rearing container for each treatment. A polystyrene cup (V-9; As One Corp., Osaka, Japan) filled with water was prepared in the rearing container. A single hole (10 mm in diameter) was made in its lid, and a paper towel was placed through the hole and touching the water to provide a watering station. Two pieces (10 cm × 15 cm) of cardboard egg carton were put into each container as shelter to mitigate cannibalism. The diet was put into a polystyrene Petri dish (90 mm diameter) placed on the egg cartons. Kimtowels (Nippon Paper Crecia Co., Ltd., Tokyo, Japan) were placed in each container to facilitate access for the crickets to the watering station and feeding area.

Cricket eggs laid in wet cotton were collected from the stock population and maintained in an incubator (MIR-554-PJ; PHC Holdings Corp., Tokyo, Japan) at 30 °C until hatching. First-instar nymphs hatched within 24 h were collected and allocated among the containers (50 in each) on day 0. The number of individuals in each container was recorded on days 6, 13, 20, 27, 34 and 41. Body weights of 10 nymphs randomly collected from each container were individually measured on an electric balance (FX-500i; A&D Co., Ltd., Tokyo, Japan) on days 20, 27, 34 and 41. When adults emerged, we calculated the developmental day and measured the body weight on an electronic balance and the head width, pronotum length, pronotum width, forewing length and hind-leg femur length with a digital calliper (CD67-S15PS; Mitsutoyo Corp., Kawasaki, Japan). Adults were removed from each container after measurement and the bioassay was continued until all surviving crickets had reached the adult stage. The bioassay consisted of three independent experimental runs.

### Calculation of population growth rate

To evaluate the daily yield, we calculated the population growth rate (PGR, g day^−1^) as:

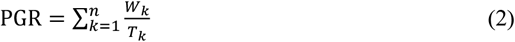

where *k =* number of crickets tested, *n* = number of adults emerged, *T*_*k*_ = developmental days from hatching to adult emergence of each individual, and *W*_*k*_ = body weight (g) of each adult within 24 h after emergence.

### Morphological analyses of mouthparts

First-instar nymphs and adults were decapitated, and the mouthparts were observed through a scanning electron microscope (VHX-DF510; Keyence Corp., Osaka, Japan). From the images, widths of the labrum sandwiched by the mandibles were calculated as an approximation of the mandibular range.

### Statistical analyses

All data analyses and visualisations were performed in R v. 4.1.0 software. Statistical differences between means of morphological data were analysed with Student’s *t*-test, Welch’s *t*-test, the Wilcoxon–Mann–Whitney *U*-test, or the Brunner–Munzel test, according to the data distribution and variance. Statistical differences of the Kaplan–Meier survival curves were analysed by log-rank test. Spearman’s rank correlation coefficient tests were performed for all correlation analyses.

## Results

### Particle size distribution

Particle sizes (mean ± SE) were 2.56 ± 0.03 mm in the granular diet and 34 ± 2 μm in the powdered diet (Figure 1). The size distributions did not overlap.

**Figure 1.**
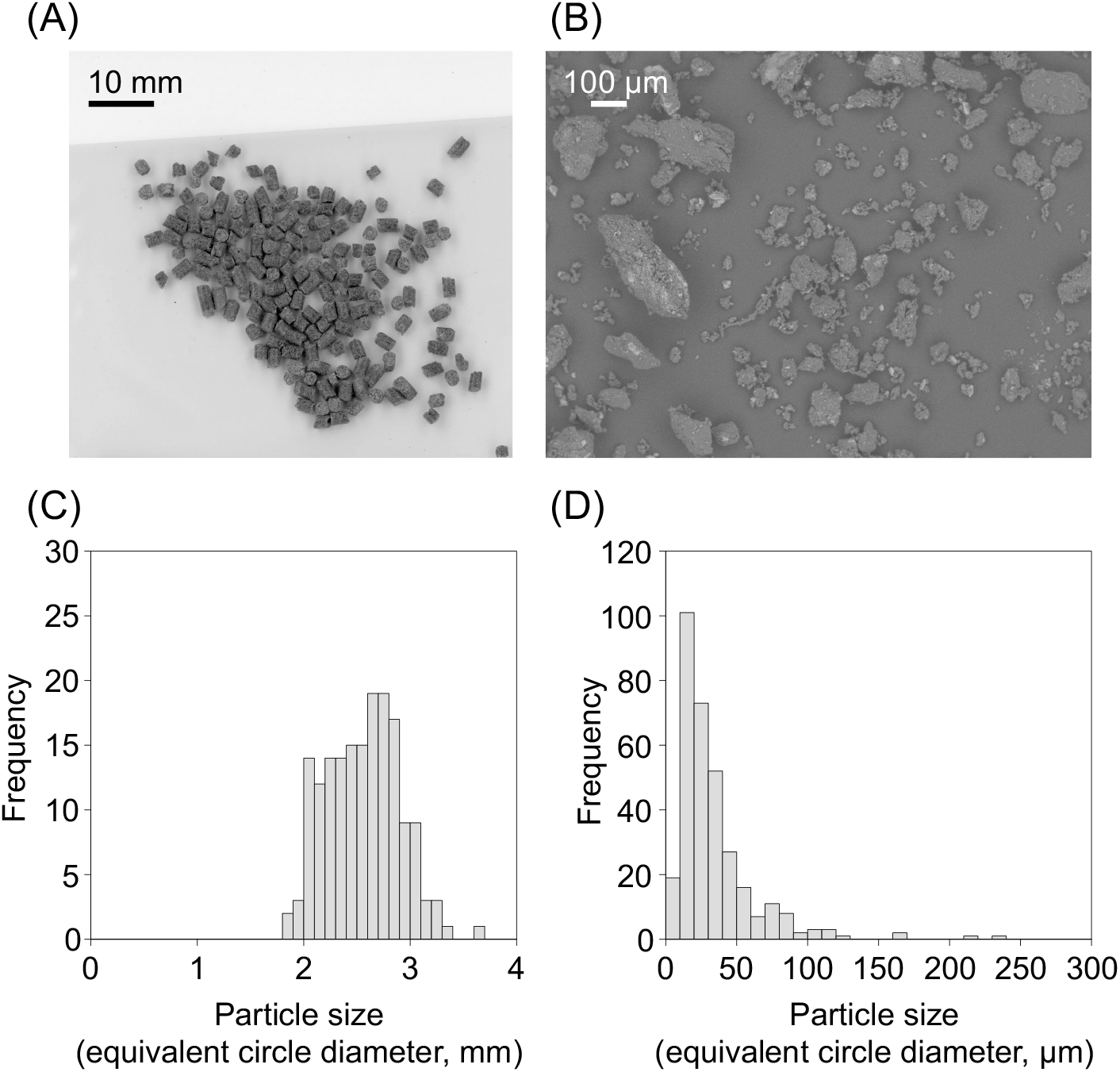
Images of (A) granular diet taken with an image scanner and (B) powdered diet taken by SEM, and particle size distributions of (C) granular and (D) powdered diet.

### Growth performances of crickets

Body weights of nymphs reared on the granular diet were significantly higher than those of nymphs reared on the powdered diet at 20, 27, 34 and 41 days after hatching (*P* < 0.05, Figure 2A). There was no significant difference in the survival curves between diets (*P* > 0.05, Figure 2B). The rate of adult emergence from nymphs fed on the granular diet was 66% ± 9% (mean ± SE)—significantly higher than that from nymphs fed on the powdered diet (39% ± 7%; *P* < 0.001, Figure 2C). The developmental time of crickets fed on the granular diet was 53 ± 1 days (mean ± SE)—significantly shorter than that of crickets fed on the powdered diet (62 ± 1 days; *P* < 0.001, Figure 2D). The adult body weight of crickets fed on the granular diet was 0.53 ± 0.01 g (mean ± SE)—significantly heavier than that of crickets fed on the powdered diet (0.38 ± 0.01 g; *P* < 0.001, Figure 2E). PGR tended to be greater in crickets fed on the granular diet (*P* = 0.09, Figure 2F). Head width, pronotum length, pronotum width, forewing length and hind-leg femur length of crickets fed on the granular diet were significantly larger than those of crickets fed on powdered diet (*P* < 0.001, Figure 3). In all adult crickets, adult body weight was correlated positively with the size of each body part (*P* < 0.001, *r* > 0.8, Figure 4A–4E) and negatively with developmental time (*P* < 0.001, *r* = −0.68, Figure 4F).

**Figure 2.**
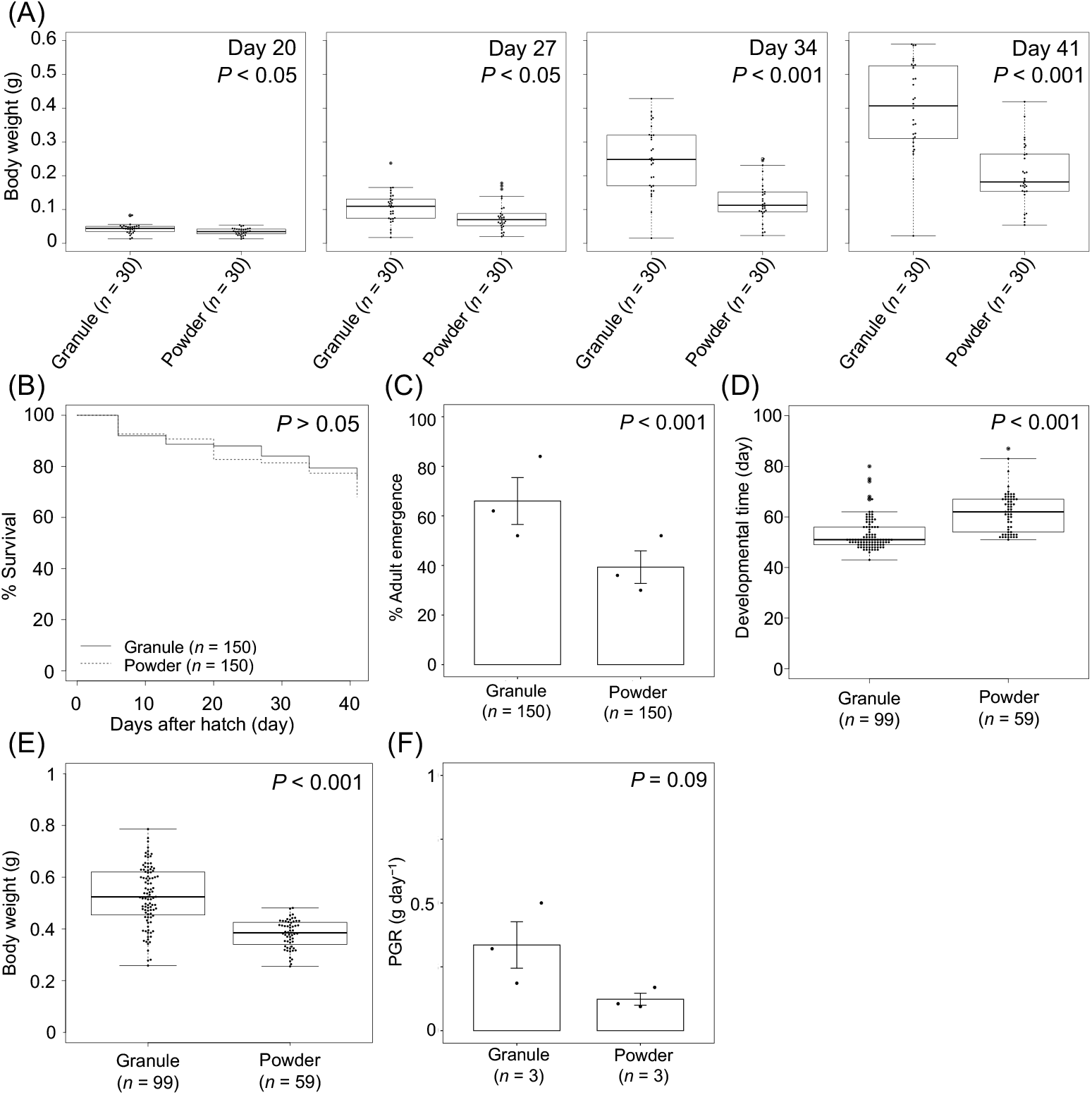
Growth performance of *Teleogryllus occipitalis* reared on a granular or a powdered diet. (A) Body weight of nymphs at 20, 27, 34 and 41 days after hatching (Wilcoxon–Mann– Whitney *U*-test for Days 20 and 27; Welch’s *t*-test for Days 34 and 41). (B) Survival curves of nymphs (log-rank test). (C) Rates of adult emergence from nymphs (Fisher’s exact test). Values are means ± SE of three independent experimental runs. (D) Developmental time of crickets (Wilcoxon–Mann–Whitney *U*-test). (E) Body weight of adult crickets (Welch’s *t*-test). (F) Population growth rate (PGR) of crickets (Student’s *t*-test). Values are means ± SE of three independent experimental runs. The initial number of crickets tested was 50 in each experimental run.

**Figure 3.**
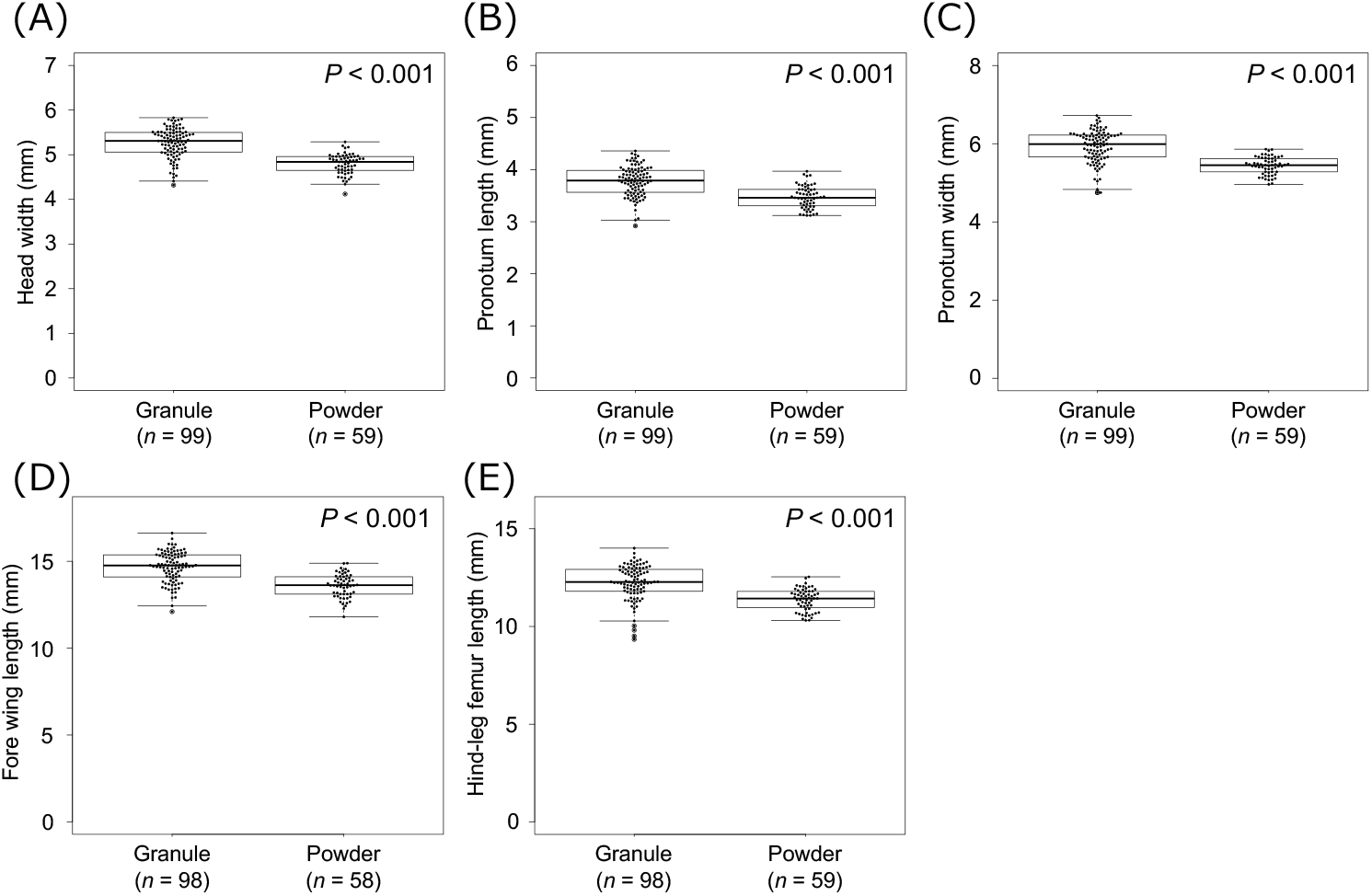
Body part sizes in adult crickets fed on a granular or a powdered diet. (A) Head width (Brunner–Munzel test). (B) Pronotum length (Welch’s *t*-test). (C) Pronotum width (Brunner–Munzel test). (D) Forewing length (Welch’s t-test). (E) Hind-leg femur length (Brunner–Munzel test).

**Figure 4.**
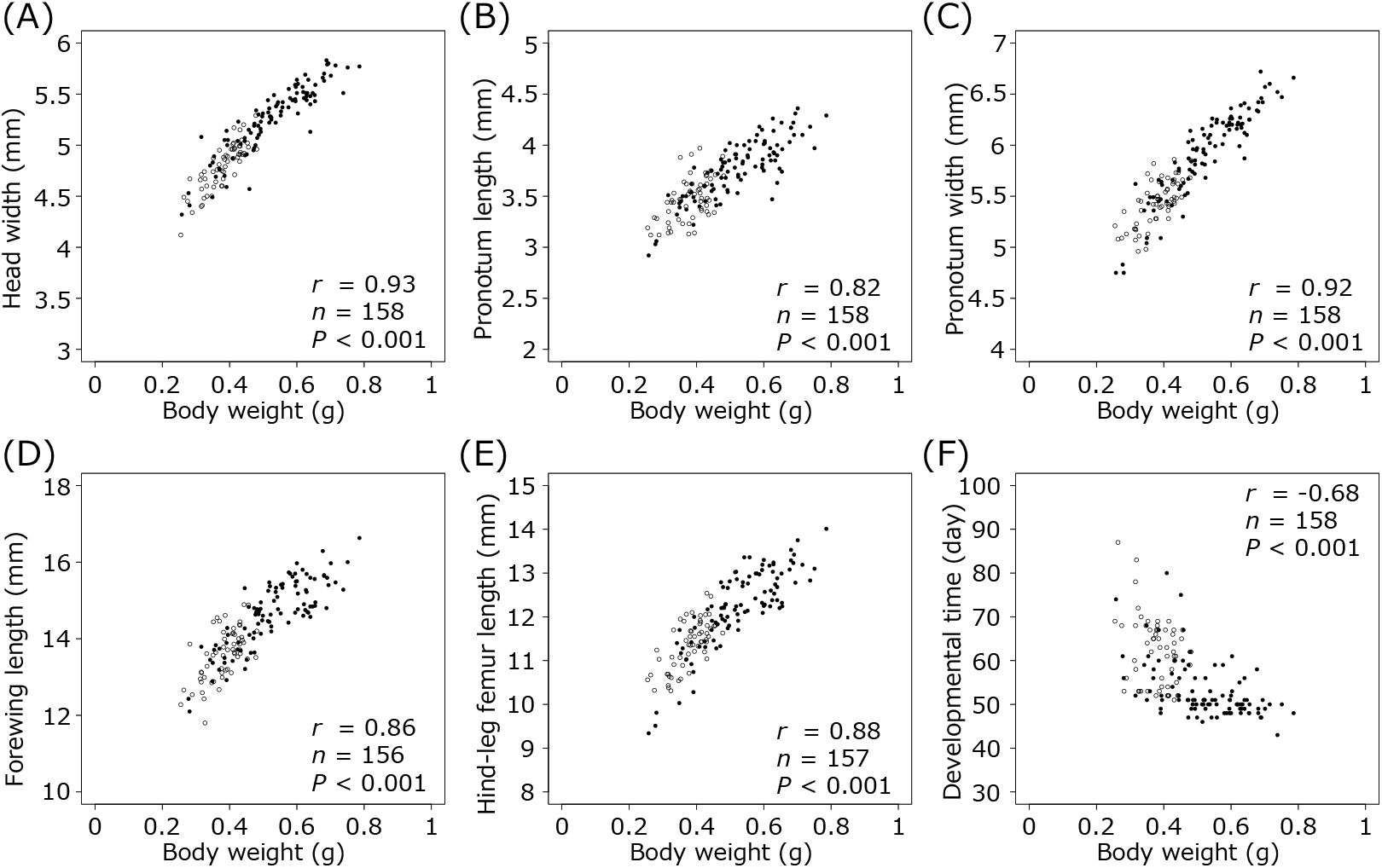
Correlations of adult body weight with (A) head width, (B) pronotum length, (C) pronotum width, (D) forewing length, (E) hind-leg femur length and (F) developmental time in *T. occipitalis* fed on ● granular or ○ powdered diet. Spearman’s rank correlation coefficient tests were performed for all correlation evaluations.

### Distance between mandibles of nymphal and adult crickets

The labrum width (mean ± SE) as an approximation of the distance between the mandibles was 0.223 ± 0.002 mm (*n* = 9) in first-instar nymphs and 1.853 ± 0.023 mm (*n* = 9) in adults (Figure 5).

**Figure 5.**
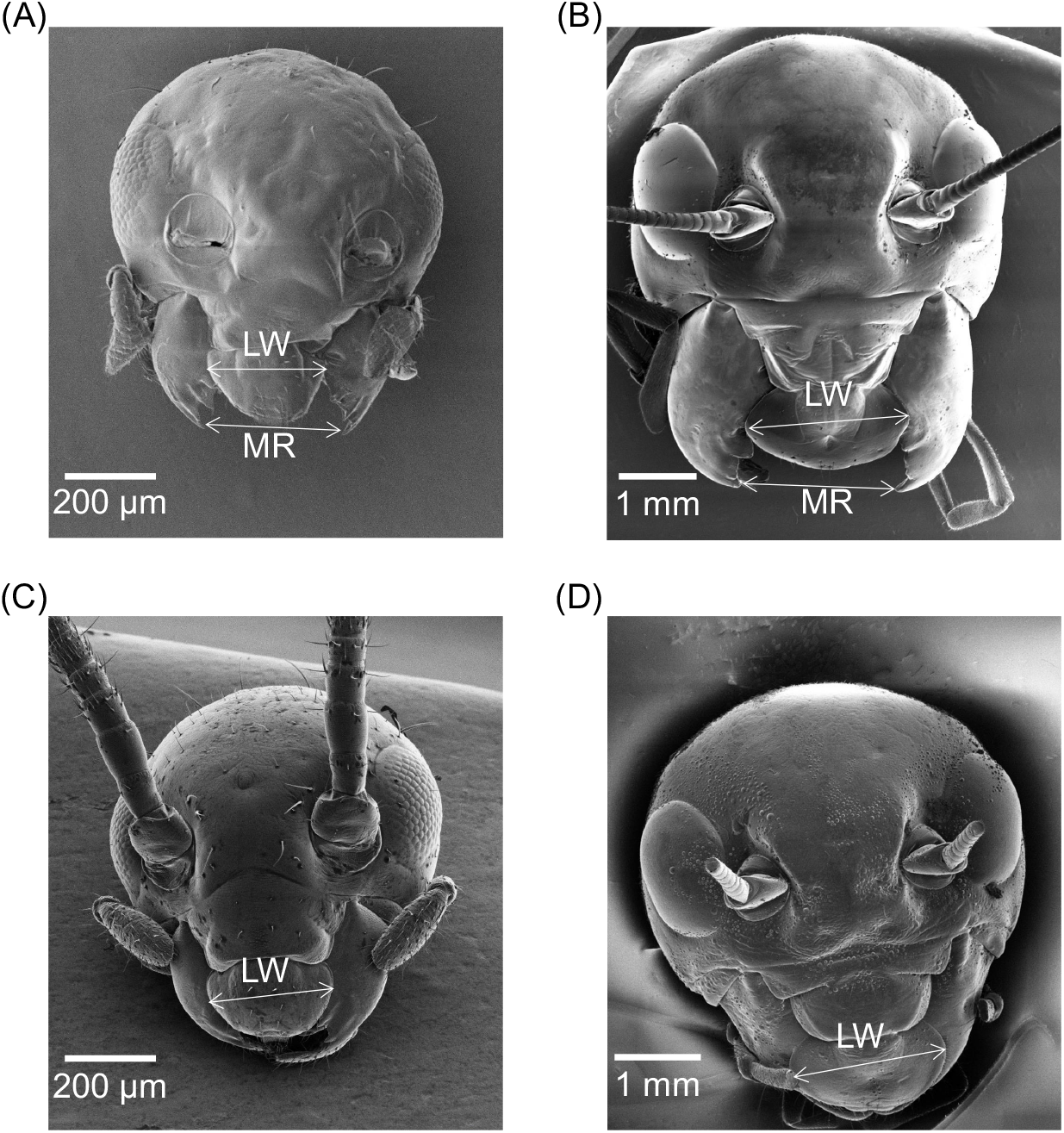
Head images of (A, C) a first-instar nymph and (B, D) an adult of *T. occipitalis* with mandibles (A, B) opened and (C, D) closed. ↔ Labrum width (LW) and mandibular range (MR).

## Discussion

As far as we know, ours is the first study to demonstrate that diet particle size affects the growth performance of crickets. Most of the growth parameters of *T. occipitalis* fed a millimetre-order granular diet were superior to those fed a micrometre-order powdered diet, even though the nutrient composition of the diets was identical.

Although the survival of crickets from hatching to adult emergence did not differ between the two diets, the body weights of nymphs and adults fed on the granular diet were higher than those of crickets fed on the powdered diet (Figure 2). However, because the body weight of insects often varies greatly depending on the timing of feeding and the water intake, the head width, pronotum length, pronotum width, forewing length and hind-leg femur length are often used as more accurate growth parameters of orthopteran insects. The lengths of all five of these body parts were significantly greater in adults that emerged from nymphs fed on the granular diet (Figure 3), and they had significant positive correlations with the adult body weight (Figure 4A–E); thus, body weight is also an accurate growth parameter, at least in *T. occipitalis* adults within a day after emergence. In contrast, adult body weight had a significant negative correlation with developmental time (Figure 4F). This result is not consistent with Masaki (1978) that head width and developmental time were positively correlated in the lawn ground cricket, *Polionemobius taprobanensis* (Walker) (Orthoptera: Trigonidiidae). In addition, there was no correlation between adult body weight and developmental time in *T. occipitalis* fed on chicken feed (unpublished data). The correlation between developmental time and adult body weight of crickets may vary with nutritional conditions during the nymphal period. Further studies are needed to investigate this hypothesis.

Mantids—predatory polyneopteran insects—grasp their prey in their forelegs. The preferred prey size of the mantid, *Hierodula crassa* Giglio-Tos (Mantodea: Mantidae), is predictive of the lengths of the femur and tibia of the foreleg and the angle between them (Holling, 1964). Crickets, which are also polyneopteran insects, grasp their food mainly in their mandibles. The estimated mandibular range in *T. occipitalis* was 0.223 mm in first-instar nymphs and 1.853 mm in adults (Figure 5). Therefore, crickets that ingested granular diet (2.56 ± 0.03 mm, Figure 1) larger than their mandibular range first had to gnaw it down to a suitable size. In contrast, the powdered diet, with a smaller particle size (34 ± 2 μm, Figure 1), was less suitable for grasping in the crickets’ mandibles, and this may have resulted in reduced food intake and smaller body size (Figures 2A, 2E, 3). Although this result seemingly contradicts the report that *A. domesticus* nymphs tend to prefer diet with small particles (Patton et al., 1967), the particle size was not indicated in that study. There may be an optimal particle size for crickets, although diet preference and growth performance may not necessarily coincide. There is also the possibility of interspecific differences between *A. domesticus* and *T. occipitalis*. Mechanoreceptors on the end of chickens’ beaks play a role in the tactile assessment of diet (Hughes and Gentle, 1995). Although insects similarly have mechanoreceptors on the galeae and maxillary palpi (Gullan and Cranston, 2010), it is unclear whether these mechanoreceptors can assess particle size. Further functional studies of the mechanoreceptors of crickets are needed for the assessment of particle size and the relationship between crickets’ diet preferences and growth performance.

Differences in physical properties dependent on particle size might also affect the growth of crickets. In general, hydrophobic powder with a particle size of 0.1 to 10 μm tends to agglomerate and float on the water surface (e.g. Zara et al., 2021). Cricket digestive systems might have difficulty digesting powdered diet owing to its lower affinity for digestive fluids. Further studies are needed to evaluate the hydrophobic properties of the two types of diets used in this study.

## Conclusion

The growth and development of *T. occipitalis* reared on a granular diet were significantly superior to those of *T. occipitalis* reared on a powdered but otherwise identical diet. This difference may be due to the presence of particles of a size that crickets’ mandibles can grasp or to size-dependent physical properties that affect affinity for digestive fluids. We plan to investigate these potential effects in order to optimize the form of diet in the production of edible crickets.

## Acknowledgements

We thank Mr Takuma Takahashi and Dr Wuled Lenggoro of the Tokyo University of Agriculture and Technology (TUAT) for useful discussions and for sharing knowledge of particle suspensions in water. We also thank Dr Susumu Inasawa of TUAT for his guidance in the use of the scanning electron microscope. This work was supported partly by the Cabinet Office, Government of Japan Cross-ministerial Moonshot Agriculture, Forestry and Fisheries Research and Development Program, “Technologies for Smart Bio-industry and Agriculture” (funded by the Bio-oriented Technology Research Advancement Institution) (JPJ009237).

## References

European Commission. 2022. Commission Implementing Regulation (EU) 2022/188 of 10 February 2022 authorising the placing on the market of frozen, dried and powder forms of Acheta domesticus as a novel food under Regulation (EU) 2015/2283 of the European Parliament and of the council, and amending Commission Implementing Regulation (EU) 2017/2470. Official Journal of the European Union, 30: 108–114. http://data.europa.eu/eli/reg_impl/2022/188/oj

Gullan, P. J. and Cranston, P. S. 2010. The insects: an outline of entomology (4th edition). Wiley-Blackwell. pp. 35.

Hanboonsong, Y., Jamjanya, T., & Durst, P. B. 2013. Edible Insect farming. http://www.fao.org/docrep/017/i3246e/i3246e00.pdf

Holling, C. S. 1964. The Analysis of Complex Population Processes. The Canadian Entomologist, 96: 335–347. 10.4039/Ent96335-1

Hughes, B. O., & Gentle, M. J. 1995. Beak trimming of poultry: Its implications for welfare. World’s Poultry Science Journal, 51: 51–61. 10.1079/WPS19950005

Kataoka, K., Minei, R., Ide, K., Ogura, A., Takeyama, H., Takeda, M., Suzuki, T., Yura, K., & Asahi, T. 2020. The draft genome dataset of the Asian cricket Teleogryllus occipitalis for molecular research toward entomophagy. Frontiers in Genetics, 11: 470. 10.3389/fgene.2020.00470

Masaki, S. 1978. Climatic adaptation and species status in the lawn ground cricket: II. Body size. Oecologia, 35(3), 343–356. 10.1007/BF00345141

Miech, P., Berggren Lindberg, J. E., Chhay, T., Khieu, B., & Jansson, A. 2016. Growth and survival of reared Cambodian field crickets (Teleogryllus testaceus) fed weeds, agricultural and food industry by-products. Journal of Insects as Food and Feed, 2: 285–292. 10.3920/JIFF2016.0028

Oonincx, D. G. A. B., van Itterbeeck, J., Heetkamp, M. J. W., van den Brand, H., van Loon, J. J. A., & van Huis, A. 2010. An exploration on greenhouse gas and ammonia production by insect species suitable for animal or human consumption. PLoS ONE, 5: 1–7. 10.1371/journal.pone.0014445

Patton, R. L. 1967. Oligidic Diets for Acheta domesticus (Orthoptera: Gryllidae). Annals of the Entomological Society of America, 60: 1238–1242. 10.1093/aesa/60.6.1238

Reece, F. N., Lott, B. D., & Deanton, J. W. 1985. The Effects of Feed Form, Grinding Method, Energy Level, and Gender on Broiler Performance in a Moderate (21 C) Environment. Poultry Science, 64: 1834–1839. 10.3382/ps.0641834

Schneider, C. A., Rasband, W. S., & Eliceiri, K. W. 2012. NIH Image to ImageJ: 25 years of image analysis. Nature Methods, 9: 671–675. 10.1038/nmeth.2089

Svihus, B., Kløvstad, K. H., Perez, V., Zimonja, O., Sahlström, S., Schüller, R. B., Jeksrud, W. K., & Prestløkken, E. 2004. Physical and nutritional effects of pelleting of broiler chicken diets made from wheat ground to different coarsenesses by the use of roller mill and hammer mill. Animal Feed Science and Technology, 117: 281–293. 10.1016/j.anifeedsci.2004.08.009

van Huis, A. 2013. Potential of insects as food and feed in assuring food security. Annual Review of Entomology, 58: 563–583. 10.1146/annurev-ento-120811-153704

van Huis, A., van Itterbeeck, J., Klunder, H., Mertens, E., Halloran, A., Muir, G. and Vantomme, P., 2013. Edible insects: future prospects for food and feed security (No. 171). Food and Agriculture Organization of the United Nations.

Zara, D. La, Zhang, F., Sun, F., Bailey, M. R., Quayle, M. J., Petersson, G., Folestad, S., & van Ommen, J. R. 2021. Drug powders with tunable wettability by atomic and molecular layer deposition: From highly hydrophilic to superhydrophobic. Applied Materials Today, 22: 1–11. 10.1016/j.apmt.2021.100945

